# Invasion blurs the biphasic mosaic, amplifies the ‘wicked problem’ through impacts on avian distributions in montane habitats

**DOI:** 10.1101/2025.04.03.646526

**Authors:** V. Jobin, C.P. Harikrishnan, A. Lele, V. Joshi, R. Chanda, S. Lawrence, P.S. Aravind, M. Mubeen, M.R. Joseph, M. Subash, R. Nandini, A. Das, D. Jathanna, VV Robin

## Abstract

Globally, managing invasive plants and habitat transformation often constitutes a ‘wicked problem’ due to highly variable impacts on threatened biodiversity. In naturally patchy habitats, invasion can blur edges, pushing species beyond natural habitats. In biphasic habitats like forest-grassland mosaics, complementary sets of specialist species offer a unique configuration to examine such impacts from both directions. In the Shola Sky Islands, extensive woody invasive stands and agriculture/production landscapes have created a set of transformed closed and open habitats structurally similar to their natural counterparts. We expect the species’ usage of such novel habitats to reflect their dietary and habitat specialisation.

We conducted 4,519 surveys across 1,204 randomly selected grid cells for the avian community, covering the global distribution of five species. For a select set of specialist and generalist species, we estimated patch occupancy and abundance using hierarchical models. We used acoustics to assess species persistence across invasion stages with automated recorders deployed across a year. We used community occurrence data to examine functional traits that correspond to habitat colonisation.

Forest species occur year-round in transformed woody habitats across all invasion stages, and habitat overstory determines avian functional diversity. Forest specialists decline in transformed habitats across high-contrast edges, while generalists increase across both high- and low-contrast edges, indicating a ‘supertramp’ strategy. Grassland specialist, however, decline strongly beyond all edges, creating a duality of losses and gains across the biphasic matrix.

We highlight the diversity of threatened species’ responses to invasion and habitat transformation, underscoring the importance of nuanced approaches to habitat restoration, particularly in biphasic habitats.

---

Large parts of the globe grapple with the impacts of novel habitats^1^ created by invasive plant species^2^. Although much attention has focused on their negative impacts^3^, these habitats pose a ‘wicked problem’^4^ for biodiversity as they may also provide services in the form of habitat or food to threatened species^5^. A common mode of landscape change in the tropics involves the expansion of woody exotics, including invasive species, triggered by the creation of production landscapes^6^. These novel landscape configurations may impact native fauna markedly but variably^7,8^. Understanding the impacts of such landscape changes, likely to be amplified by climate change, can help us manage and conserve biodiversity.

Woody plant invasion can ‘blur’ edges of naturally patchy habitats and create a low-contrast matrix adjacent to habitat patches such as rainforests. Resultant semi-natural matrices with an altered gradient of habitat structure and microclimate^9^ can result in continuous faunal immigration from natural habitat patches. Such colonisation of novel habitats will alter species distributions^10^ and probabilities of occupancy, dependent on the proximity to natural patches^11^. Since habitat-generalist species are prone to use a greater variety of habitats in the landscape, they should be less affected by the contrast of the surrounding matrix than habitat-specialist species^12^.

Naturally occurring high-contrast mosaics of dissimilar habitats (e.g., bi-phasic mosaics) in different parts of the world, with complementary sets of specialised species, offer a unique opportunity to study the two sides of the impacts of landscape changes. In bi-phasic forest-grassland mosaics, both habitats with their specialist and generalist species occur in the same climatic zone (montane ecosystems). Species may occupy habitats based on vegetation structure and composition^13,14^, with their occurrences explained by their dietary specialisation^15^ or edge/interior preferences^16^. When such habitats are modified due to production landscapes or through woody invasion, a semi-natural matrix with ‘softened’ edges is created^17^. Studies across multiple climatic zones either highlight the advantage forest specialists have over grassland specialists in high land-use intensity^18^ or show no differential responses between specialists of grassland or forests to land cover change^19^. Responses to such modifications within the same climatic zone, studied from the perspective of both sets of species of the contrasting habitats - the open and wooded habitats - have yet to be investigated.

The Shola Sky Islands (SSI) of the Western Ghats are a naturally bi-phasic mosaic of montane forests and grasslands^20^. These fragmented habitats host several endemic species, including taxa unique to each sky island, as well as both forest and grassland specialists. Over the last century, the invasion of timber trees (*Acacia mearnsii, Eucalyptus spp*., *Pinus spp*.) and the spread of human-modified habitats into natural grasslands have transformed 60% of the landscape^21^. The transformed mosaics - wooded (timber stands caused by planting and invasion) vs open habitats (tea plantations, agriculture and human settlements), can approximate the vegetation structure of natural mosaics - closed canopy (shola forests/ natural wooded habitat) vs open (shola grasslands / natural open habitats). A few bird species of SSI are known to occupy most habitats (habitat-generalist)^22^ - natural and transformed, but the influences on specialist species in each habitat configuration are unknown and are likely to provide insights into impacts that may be further exacerbated by climate change – a likely increase in wooded areas^23^.

This study examines bird occurrence and abundance across natural and transformed high-contrast habitats. Additionally, we ask if transformed habitats – whether wooded or open – play a transient role as habitat patches relative to natural habitats. We expect forest specialist species, in particular, to show reduced occupancy and abundance in transformed habitats compared to generalist species.

We surveyed two of the largest sky islands above 1,400m ASL, following land cover classes: forests, timber plantations, tea plantations, settlements, and agriculture, based on Arasumani et al. (2018)^21^. We randomly selected grid cells covering 0.5% of the landscape and managed to sample 747 grid cells of 1ha area and 305 grids of 4ha area. We sampled a grassland specialist in a total of 899 grid cells^24^(Please see *Detailed methods*). Our sampling included three levels – a) targeted species (occupancy and density), b) avian community (presence/absence) across the study grids, and c) long-term persistence of target species over a year in select habitats. The target species included eight forest bird taxa (six habitat specialists) and one specialist grassland bird (Figure 1 (D)). We surveyed each grid cell for forest birds using playback songs between June 2019 and May 2023, with a maximum of four visits per cell. The occurrence of the grassland bird was also recorded. We included the survey data^24^ of the grassland bird collected between October 2017 and May 2018 for the analyses. Altogether, in our study of nine species, we covered the global distribution of five endemic endangered birds through 3,365 surveys of 899 sites. We also collected site-level variables for landscape, topography, climate, vegetation structure and composition. Visit-level covariates (wind and observer experience) were also recorded during field surveys.

**Figure 1:**
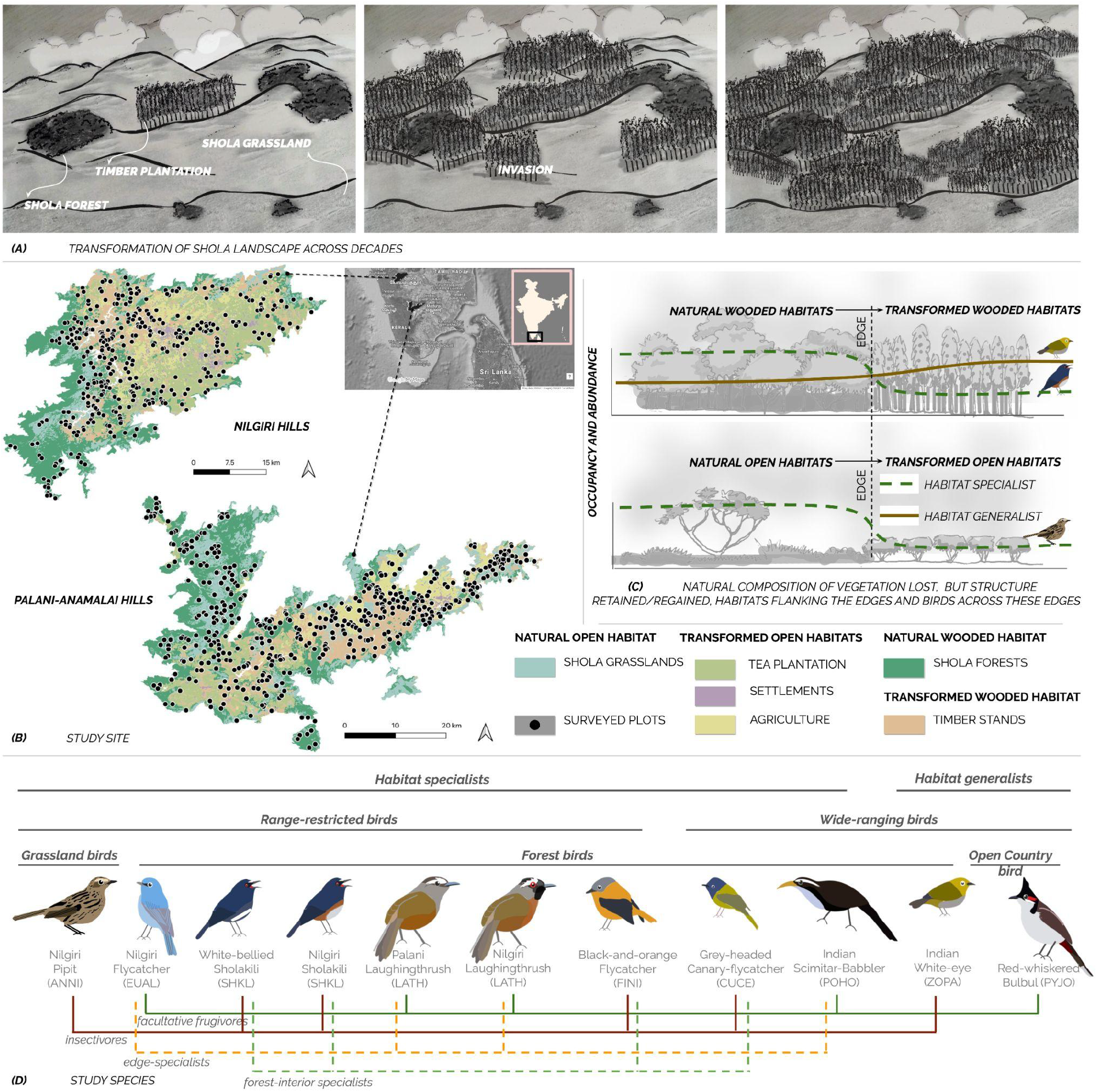
(A): Illustration of the landscape change across the biphasic matrix across decades. (B) Maps showing the biphasic landscape with natural wooded habitats (*shola* forests), natural open habitats (*shola* grasslands), and their transformed counterparts – transformed wooded habitats (timber stands caused by planting and invasion) and transformed open habitats – tea plantations, agriculture and settlements; the dots indicate the distribution of survey sites. (C) We predict a decline (with distance) in specialist and generalist birds transitioning from (i) natural wooded habitats to transformed wooded habitats and (ii) natural open habitats to transformed open habitats. (D) Study species, categorised based on their habitats, ranges and diet.

We employed a two-step modelling procedure^25^ for both occupancy and abundance analyses in *Unmarked*^*26*^. For occupancy analyses of the grassland specialist, we modelled the site area as a covariate for species-level detection and occupancy. Additionally, for abundance analyses of forest species, we first used offset(log(Area)) as an offset ^27^ to control for variable site area. We conducted **χ**^2^goodness-of-fit tests and constructed response curves for each covariate using model-averaged predictions with 95% confidence intervals^28^. To assess the impact of altered land cover on the avian community, we also recorded the occurrences of all avian taxa within woody sites (including study taxa) and used these to estimate avian functional diversity based on Bregman et al.^29^.

To examine the year-round occurrence of species in transformed wooded habitats, we deployed Automatic Recording Units (ARU) for 12 months from July 2018 to June 2019 at three sites (each >500 m apart) across a successional gradient of woodland transformation.

The naive occupancy of each taxon within natural habitats and the transformed habitats was comparable, except for the grassland specialist bird. The occupancy of study birds is greatly affected by climatic variables like precipitation seasonality, mean temperature of the coldest quarter and distance from native habitats. The structural variables of vegetation show no significant effect on their occupancy (Supplementary Table 1).

On the other hand, in abundance models, patterns of density changed based on the diet-specialisation of the birds. Invertivores show strong relationships with vegetation variables, notably structural variables like foliage density of the understory. Facultative frugivores show marked variation across the presence of different fruiting plant species in the understory (e.g. *Daphniphyllum neilgherrense, Rubus spp*.). Forest specialist birds correlate positively with forest-specific vegetation variables like the basal area and canopy cover. Range-restricted birds show an apparent increase in density with landscape variables, particularly elevation and proximity to forest. However, unlike the typical expectation, habitat specialist birds in this landscape were not restricted to the interiors of the native habitats - natural open or wooded habitats. The distance from the edges impacted the forest birds’ occupancy outside their native habitat. The forest specialists showed a marked decrease in occurrence outside any wooded habitats - natural or transformed. However, they show a gradual reduction in their abundance across the edges between natural wooded and transformed wooded habitats. The grassland specialist bird, however, has a similar, lowered occurrence pattern across both low- and high-contrast edges (Fig 2A).

**Figure 2:**
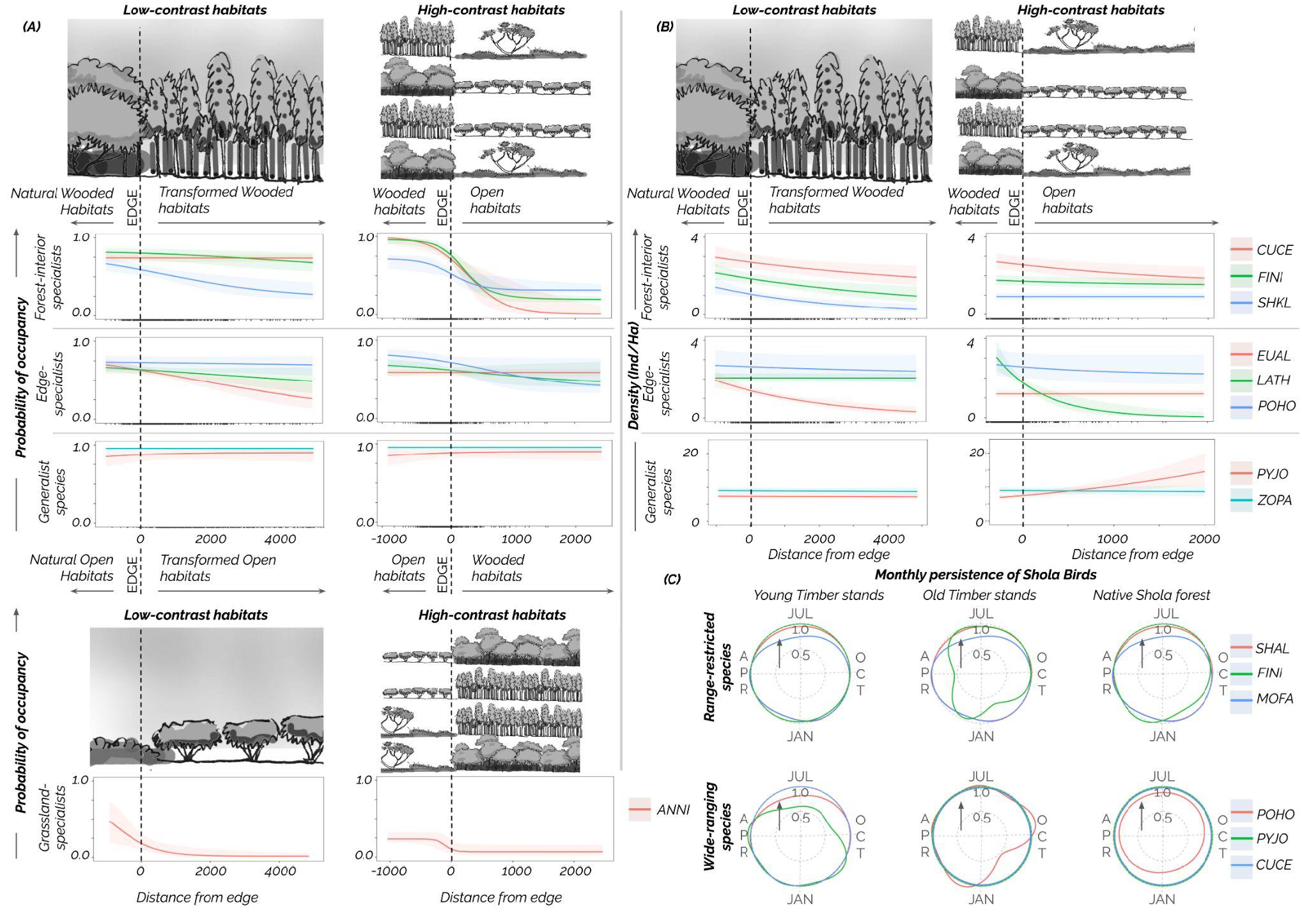
(A) Forest specialists occupy transformed woodlands to a greater extent when these habitats are adjacent to the natural woodlands (low-contrast habitats) than to high-contrast habitats, open habitats like grasslands or agriculture; this pattern is reduced in edge-specialist birds and but not in the generalists that occupy all transformed and natural habitats. The grassland specialist experiences a sharp decline in occupancy from natural open habitats to transformed low-contrast habitats and a stronger reduction in high-contrast woodland habitats. (B) The abundance of forest interior specialists and edge specialists gradually declines from natural to transformed woodlands and further to high-contrast open habitats. The abundance of generalist woodland species rises from woodlands to high-contrast open habitats. (C) Automatic Recording Units reveal the year-round occurrence (probability) of range-restricted and wide-ranging species of forest birds in transformed woodlands at all stages of invasion.

Acoustic data supports the occurrence of forest birds throughout the year in the transformed woodlands. The specialist and generalist birds show similar trends across all months (Fig 2C).

The observed functional diversity was significantly higher than the null expectation for both invertivores and facultative frugivores in Shola forests. However, for stands of non-native trees, the observed functional diversity varied based on the overstory tree species, most likely as they affect understory composition^30,31^ (Supplementary Figure 1). All statistical results are from two-tailed Wilcoxon signed-ranks tests.

Forest birds, specialists or generalists, occur in the transformed wooded habitats throughout the year, suggesting a lack of transient movement between natural and transformed habitats, or at least the persistence of such forays through the year. However, the distance of occurrence from the edge of forests was far greater than several times the home range dimensions of these restricted birds. This suggests that these are not short-distance foraging forays into the novel habitats from the forest edges but territory-holding and breeding, possibly maintained through source-sink dynamics^32,33^ between forests and transformed wooded habitats.

Forest specialists show little difference in their preferences for transformed wooded habitats when compared to their natural habitats and no edge-avoidance for edges between high-contrasting habitats but only gradual declines in occurrence^34,35^ across low-contrasting habitats. On the other hand, grassland specialists indicate high edge avoidance in both high- and low-contrasting edges, showing their avoidance of wooded habitats altogether. The ongoing invasion may thus impact their occurrence in the landscape. Generalist species^36^, on the other hand, are consistent across most habitats, with their numbers increasing in open habitats, resulting in greater competition for grassland specialists. Such generalist species appear to follow a ‘supertramp’ strategy^36^ with their ability to occupy a diversity of habitats in these species-poor mountain-top islands^20,37^.

Taken together, the human-led modification of the natural high-contrast habitat matrix of forests and grasslands into transformed woodlands and open habitats does not impact all birds equally or even in the same direction. The transformed woodlands, both planted and invaded, have facilitated most forest birds, with ‘supertramps’ exploiting the novel habitats to typify a ‘wicked problem’. Nonetheless, the grassland specialist has declined precipitously-limited by the expansion of invasive woody trees along with agricultural and human-use landscapes into the natural grasslands. Climate change^38^ is projected to reduce grassland habitats further, while forest birds may see future expansion. Our study highlights a wicked problem posed by a landscape-scale invasion, hastening the loss of a species while facilitating the spread of several others.

## Supporting information

Detailed methods

