## Supplementary material for "Invasion blurs the biphasic mosaic, amplifies the ‘wicked problem’ through impacts on avian distributions in montane habitats": Detailed methods

***Year-round occurrence of birds in the landscape***

Since the forays of forest birds between natural wooded habitats and transformed wooded habitats can affect our results[1], we verified the difference in the year-round occurrence of birds in both habitats. We used acoustically collected occurrence data across a single year [2] to reveal patterns of occurrence of shola bird biodiversity in transformed wooded habitats. In one sky island, we deployed automated recorder units (ARUs) in three habitats: young timber stands (new invasion), old timber stands, and natural woodlands - shola forests, between July 2018 and June 2019. After subsampling the data for morning hours (6 am to 9 am) every alternate day, we manually annotated the calls using Kaleidoscope Pro[3]. We modeled the monthly occupancy patterns of range-restricted (*Montecincla fairbanki*, *Sholicola albiventris*, *Ficedula nigrorufa*) and wide-ranging (*Culicicapa ceylonensis*, *Pycnonotus jocosus*, *Pomatorhinus horsfieldii*) bird species using Generalized Additive Modeling[4]. This analysis included ~600 hours of data, with 282725 detections across the habitat types.

***Occupancy and Abundance of birds in the landscape***

We surveyed areas above 1400m MSL within the two largest Shola sky islands, focusing on the following land cover types: natural woodlands - Shola forests, transformed woodlands - timber stands (invasion and plantations), and natural open habitats - Shola grasslands, along with the transformed open habitats – tea plantations, settlements, and agriculture, based on Arasumani et al. (2018)[5]. We created grid points spaced 100m apart across the area and selected 0.5% using ArcGIS. We placed nested square cells of 1ha and 4ha over the selected points. We surveyed six forest bird species within the 1ha cells and two forest bird species within the 4ha cells based on the expected home range sizes.

In all grids, we recorded all birds detected aurally and visually, with all data collectors trained to recognise the community of local birds (details further below).

We selected common resident forest understory birds with small home ranges - Sholakilis [White-bellied Sholakili (*Sholicola albiventris*) and Nilgiri Sholakili (*Sholicola major*)], Laughingthrushes [Palani Laughingthrush (*Montecincla fairbanki*) and Nilgiri Laughingthrush (*Montecincla cachinnans*)], Black-and-orange Flycatcher (*Ficedula nigrorufa*), Nilgiri Flycatcher (*Eumyias albicaudatus*), Gray-headed Canary-flycatcher (*Culicicapa ceylonensis*), Indian White-eye (*Zosterops palpebrosus*), Red-whiskered Bulbul (*Pycnonotus jocosus*), and Indian Scimitar-babbler (*Pomatorhinus horsfieldii*). The first six taxa had a rough home range of ~1ha[6,7]. The latter two species have a moderately larger home range (~4ha). For Red-whiskered Bulbul, the distance is calculated based on their median displacements during the median gut passage times for seeds (130m)[8]. Shankar Raman[9] assessed the density of the Indian Scimitar-Babbler through spot-mapping technique as 2 individuals/ 4ha. We also analysed a grassland bird species, Nilgiri Pipit (*Anthus nilghiriensis*), using data from Lele et al. (2020)[10]. In the wooded habitats (natural and transformed), we selected 750 potential sites and managed to survey 599 sites. Within the transformed open habitats, we selected 530 sites and surveyed 148, while the data for grassland specialist bird is from 152 sites surveyed from a selected 202 potential sites. Due to inaccessibility and recent land conversion into agricultural fields, our analysis data is restricted to 747 grid cells of 1ha area (299 cells nested within the 4ha grid cells) and 305 4ha grid cells. We surveyed the grassland bird Nilgiri Pipit, in the wooded and transformed open habitats, as well, using similar methods to those of Lele et al. (2020)[10]. The final number of surveyed sites for grassland birds is 899.

We examined the set of variables responsible for the patterns of distribution and abundance of our study birds across the mountain tops: landscape variables, topographical variables, climate variables, physiognomic variables and floristic variables (Supplementary Table 2). The latter two sets were collected on the ground within the survey cells. We employed the single-season occupancy model of Mackenzie et al. (2002)[11] to determine the factors driving the distribution and the *N*-mixture model of Royle (2004)[12] to investigate the drivers of abundance of the study birds through multiple non-mutually exclusive hypotheses based on the available literature and expert opinions. Visit-level covariates (wind and observer experience) were also recorded in the field. To disentangle ecological processes responsible for the distribution and abundance of our study birds from the process affecting the detection (species-level for occupancy, individual-level for *N*-mixture) of these birds, we surveyed each site 1-4 times using playback songs of our study species between June 2019 and May 2023, with a maximum of four visits per site. The average survey duration for 1ha is 10.27 (+2.52, SE) minutes, and for 4ha is 23.98 (+7.74, SE) minutes.

We examined the covariates, and within the sets of covariates for each species, we eliminated the covariates with high collinearity (Pearson's $|r| > 0.7$)[13]. We employed a two-step modelling procedure[14] for both occupancy and abundance analyses, performed using the package *Unmarked*[15] in the statistical software R (R Core Team, 2018). First, we used a universal covariate structure to model the variation in occupancy or abundance while exploring various combinations of covariates for the observation processes that influence detection. The models were based on sets of covariates that affect visual (vegetation structure like canopy and leaf density) and aural cues (wind speed with vegetation structure) to bird detection and covariates particular to the native habitats of these birds - the distance from shola forest or wooded areas, moss cover, and basal area of shola trees. In the second step, we selected the covariate combination of the best-supported model with different covariate combinations for occupancy or abundance to determine the final most supported model for occupancy and *N*-mixture analyses, respectively[16]. The sets of covariates were categorised as covariates specific to Shola forests, landscape variables, and a combination of local scale vegetation variables and landscape variables. In the analysis for each taxon, the number of covariates used was thirteen or less, and the number of models examined was less than the sample size, based on Burnham and Anderson[17]. For Nilgiri Pipit, the site area was used as a covariate to control for species detection and occupancy. For the abundance of the forest birds, the log of the site area was used as an offset to control for site area. The adequacy of the model was determined by the chi-square goodness-of-fit test with 10,000 (for occupancy models) and 99 (for abundance models) parametric bootstrap simulations on the model with the maximum number of parameters[18]. Response curves for each covariate were constructed using model-averaged predictions with a confidence interval of 95%[17]. When the $\chi^2$ goodness-of-fit test indicated moderate over-dispersion of latent abundances relative to the model (P-value < 0.05), the estimated overdispersion factor ($\hat{c}$) was used to derive QAIC values for model selection and to adjust estimated variances.

***Avian functional traits across wooded habitats in the Shola landscape***

Apart from the subset of focussed study species, the occurrences of other birds within the wooded sites were also noted. The community data was used to measure the avian functional diversity across various wooded habitats in the Shola landscape based on Bregman et al. (2016)[19] to assess the impact of altered land cover on the avian community.

We extracted seven morphometric measurements from the AVONET database[20] of all the birds in our study sites (54 species): beak length, width and depth, wing length, tarsus length and tail length. To remove the correlation between the functional traits of the birds, particularly body size with other traits, we use ordination techniques to procure three trait axes: locomotory traits (PCA between tarsus vs tail/wing ratio), trophic traits (PCA between beak morphometry: beak length, beak depth, beak width), and size traits (PCA between first PCA components of the former two PCAs). We calculated Functional Distance (FD) by converting the trait matrix of all birds into a distance matrix, with which we produced a dendrogram using Euclidean distance and UPGMA clustering[21]. The FD of a given community (species pool sampled in a given grid cell) is the total branch length connecting all the species in the community. We assessed how the raw FD of communities varied across land covers. We compared them with the FDs of communities within the land cover based on null expectations drawn from 999 random communities (species richness was the same as the observed, and the probability of the presence of each species was determined by its frequency across all communities). Observed values of standardized FD were pooled for each land-cover category. All statistical results are from two-tailed Wilcoxon signed-ranks tests.
