## Supplementary Table 1 for "Invasion blurs the biphasic mosaic, amplifies the ‘wicked problem’ through impacts on avian distributions in montane habitats"

Supplementary Table 1: The table shows model-averaged estimates of all covariates from the occupancy and abundance analyses carried out for each species. Species - SHKL: *Sholicola albiventris* and *Sholicola major*; LATH: *Montecincla fairbanki* and *Montecincla cachinnans*; FINI: *Ficedula nigrorufa*; EUAL: *Eumyias albicaudatus*; CUCE: *Culicicapa ceylonensis*; ZOPA: *Zosterops palpebrosus*; PYJO: *Pycnonotus jocosus*; POHO: *Pomatorhinus horsfieldii*; and ANNI: *Anthus nilghiriensis*. □ is the model-averaged beta coefficient, and SE is the standard error. Psi (Ψ) is the probability of occupancy of a species. Lambda (λ) is the estimated latent abundance of a species in a site. The probability of occupancy for most study species was correlated with landscape-level covariates (Elevation, Presence of Shola forest, Wetness Index, etc.). The probability of occupancy was not correlated with site-level variables, namely the vegetation variables like the canopy cover, basal area and foliage density. The estimates coloured in shades of red show negative estimates, and shades of green show positive estimates. Estimates in yellow are closer to zero.

| | **Model-averaged estimates of □-coefficients from abundance analyses** | | | | | | | | | | | | | | | | | | |
|---|---|---|---|---|---|---|---|---|---|---|---|---|---|---|---|---|---|---|---|
| | **Variables** | **CUCE** | | **EUAL** | | **FINI** | | **LATH** | | **POHO** | | **PYJO** | | **SHKL** | | **ZOPA** | | **ANNI** | |
| | | □ | SE | □ | SE | □ | SE | □ | SE | □ | SE | □ | SE | □ | SE | □ | SE | □ | SE |
| **Structural** | Canopy cover | 0.5 | 0.1 | 0.4 | 0.0 | | | 0.2 | 0.1 | 0.5 | 0.1 | -0.9 | 0.1 | 0.5 | 0.0 | 0.2 | 0.0 | | |
| | Ferns | 0.1 | 0.0 | | | 0.2 | 0.0 | | | | | | | | | 0.2 | 0.0 | | |
| | Leaf Litter | | | 0.2 | 0.0 | | | 0.0 | 0.0 | | | | | | | | | | |
| | Humidity (moss cover as a proxy) | 0.1 | 0.0 | 0.0 | 0.0 | 0.3 | 0.1 | 0.1 | 0.0 | 0.0 | 0.0 | | | 0.2 | 0.0 | -0.2 | 0.0 | | |
| | Total_basal_area | 0.0 | 0.0 | 0.0 | 0.0 | 0.4 | 0.1 | 0.0 | 0.0 | 0.2 | 0.2 | 0.1 | 0.0 | | | -0.1 | 0.0 | | |
| | Foliage density b/w 0-2 m | 0.1 | 0.0 | 0.2 | 0.0 | 0.3 | 0.0 | | | | | | | | | | | | |
| | Foliage density b/w 0-3 m | | | | | | | | | | | | | 0.2 | 0.0 | | | | |
| | Foliage density b/w 0-5 m | | | | | | | 0.3 | 0.0 | 0.1 | 0.1 | 0.1 | 0.0 | | | | | | |
| | Foliage density b/w 2-5 m | 0.0 | 0.0 | 0.1 | 0.1 | | | | | | | | | | | 0.0 | 0.0 | | |
| **Compositional** | Bamboo | | | | | 0.2 | 0.1 | | | 0.0 | 0.0 | | | | | | | | |
| | *Daphniphyllum* | | | 0.0 | 0.0 | | | 0.0 | 0.0 | 0.3 | 0.2 | 0.3 | 0.0 | | | | | | |
| | *Lantana* | | | | | | | | | | | 0.2 | 0.0 | | | | | | |
| | *Rubus* | | | 0.1 | 0.0 | | | 0.1 | 0.0 | 0.0 | 0.0 | 0.2 | 0.0 | | | | | | |
| | Basal area of Shola trees | 0.0 | 0.0 | 0.0 | 0.0 | 0.0 | 0.1 | 0.1 | 0.0 | 0.1 | 0.0 | | | | | | | | |
| **Climatic** | Mean Temperature of Coldest Qtr | | | -0.2 | 0.0 | -0.3 | 0.1 | | | | | | | -0.2 | 0.0 | | | | |
| | Precipitation seasonality | 0.0 | 0.0 | | | | | -0.2 | 0.0 | | | | | -0.1 | 0.0 | 0.0 | 0.0 | | |

| | | | | | | | | | | | | | | | | | |
|---|---|---|---|---|---|---|---|---|---|---|---|---|---|---|---|---|---|
| | Mean Precipitation of Warmest Qtr | | | 0.1 | 0.0 | | | | | | | | | | | | |
| | Temp. Seasonality | | | | | | | -0.3 | 0.0 | -0.3 | 0.0 | 0.3 | 0.0 | | | 0.0 | 0.0 |
| **Landscape** | Elevation | -0.1 | 0.0 | | | | | | | | | 0.0 | 0.0 | | | | |
| | Open human-modified area | | | | | | | | | | | 0.0 | 0.0 | | | | |
| | Productive area | -0.3 | 0.0 | 0.0 | 0.1 | | | -0.1 | 0.1 | | | | | -0.6 | 0.0 | 0.1 | 0.0 |
| | Distance from Shola edges | -0.2 | 0.1 | -0.3 | 0.0 | -0.2 | 0.0 | -0.1 | 0.0 | -0.1 | 0.0 | 0.0 | 0.0 | -0.3 | 0.0 | 0.0 | 0.0 |
| | Distance from Wooded habitat edges | -0.1 | 0.0 | 0.0 | 0.0 | -0.2 | 0.1 | -0.3 | 0.3 | -0.2 | 0.1 | 0.2 | 0.0 | -0.1 | 0.0 | 0.0 | 0.0 |
| | Proximity to stream (TWI) | 0.0 | 0.0 | -0.1 | 0.0 | 0.0 | 0.0 | -0.1 | 0.0 | | | | | 0.0 | 0.0 | | |

**Model-averaged estimates of covariates from occupancy analyses**

| | **Variables** | **CUCE** | | **EUAL** | | **FINI** | | **LATH** | | **POHO** | | **PYJO** | | **SHKL** | | **ZOPA** | | **ANNI** | |
|---|---|---|---|---|---|---|---|---|---|---|---|---|---|---|---|---|---|---|---|
| | | □ | SE | □ | SE | □ | SE | □ | SE | □ | SE | □ | SE | □ | SE | □ | SE | □ | SE |
| **Structural** | Canopy cover | 0.8 | 0.2 | 0.5 | 0.1 | | | 0.6 | 0.1 | 0.6 | 0.2 | -1.5 | 0.2 | 0.7 | 0.2 | 0.9 | 0.1 | | |
| | Ferns | 0.5 | 0.2 | | | 0.5 | 0.2 | | | | | | | | | 0.4 | 0.2 | | |
| | Leaf Litter | | | 0.3 | 0.1 | | | 0.2 | 0.1 | | | | | | | | | | |
| | Humidity (moss cover as a proxy) | 0.8 | 0.3 | 0.1 | 0.1 | 1.6 | 0.3 | 0.4 | 0.1 | 1.0 | 0.3 | | | 0.9 | 0.3 | 1.2 | 0.5 | | |
| | Total_basal_area | 0.4 | 0.4 | 0.0 | 0.1 | 0.8 | 0.4 | 0.0 | 0.1 | 0.8 | 0.4 | -0.3 | 0.2 | | | 0.0 | 0.2 | | |
| | Foliage density b/w 0-2 m | 0.1 | 0.1 | 0.3 | 0.0 | 0.5 | 0.1 | | | | | | | | | | | | |
| | Foliage density b/w 0-3 m | | | | | | | | | | | | | 0.5 | 0.1 | | | | |
| | Foliage density b/w 0-5 m | | | | | | | 0.6 | 0.1 | 0.4 | 0.1 | 0.0 | 0.1 | | | | | | |
| | Foliage density b/w 2-5 m | 0.4 | 0.3 | 0.4 | 0.1 | | | | | | | | | | | 0.5 | 0.4 | | |
| **Compositional** | Bamboo | | | | | 2.1 | 3.3 | | | 0.1 | 0.1 | | | | | | | | |
| | *Daphniphyllum* | | | 0.0 | 0.0 | | | 0.3 | 0.0 | 0.1 | 0.5 | 0.9 | 0.3 | | | | | | |
| | *Lantana* | | | | | | | | | | | 1.4 | 0.4 | | | | | | |
| | *Rubus* | | | 0.4 | 0.0 | | | 0.3 | 0.0 | 0.1 | 0.0 | 0.8 | 0.1 | | | | | | |
| | Basal area of Shola trees | 0.9 | 0.4 | 0.1 | 0.1 | 0.0 | 0.2 | 0.0 | 0.1 | 0.7 | 0.3 | | | | | | | | |
| **Climatic** | Mean Temperature of Coldest Qtr | | | -0.2 | 0.0 | -0.5 | 0.1 | | | | | | | -0.4 | 0.1 | | | -1.8 | 0.7 |

| | | | | | | | | | | | | | | | | | | | |
|---|---|---|---|---|---|---|---|---|---|---|---|---|---|---|---|---|---|---|---|
| | Precipitation seasonality | -0.1 | 0.1 | | | | | -0.4 | 0.1 | | | | | -0.2 | 0.1 | 0.1 | 0.1 | 0.8 | 0.9 |
| | Mean Precipitation of Warmest Qtr | | | 0.1 | 0.0 | | | | | | | | | | | | | 0.5 | 1.0 |
| | Temp. Seasonality | | | | | | | -0.5 | 0.1 | -0.4 | 0.1 | 0.7 | 0.2 | | | -0.1 | 0.1 | -1.2 | 0.6 |
| **Landscape** | Elevation | -0.2 | 0.1 | | | | | | | | | -0.1 | 0.2 | | | | | | |
| | Open human-modified area | | | | | | | | | | | -0.1 | 0.0 | | | | | | |
| | Productive area | -0.6 | 0.2 | -0.3 | 0.1 | | | -0.4 | 0.2 | | | | | -0.8 | 0.2 | -0.2 | 0.2 | | |
| | Distance from Shola edges | -0.5 | 0.1 | -0.4 | 0.1 | -0.5 | 0.1 | -0.3 | 0.1 | -0.3 | 0.1 | 0.3 | 0.3 | -0.7 | 0.2 | -0.1 | 0.1 | | |
| | Distance from Wooded habitat edges | -1.4 | 0.3 | -0.3 | 0.1 | -1.0 | 0.4 | -1.0 | 0.2 | -0.5 | 0.2 | 1.0 | 0.4 | -1.6 | 0.5 | -0.1 | 0.1 | | |
| | Distance from Grassland edges | | | | | | | | | | | | | | | | | -1.9 | 0.3 |
| | Distance from Open habitat edges (transformed) | | | | | | | | | | | | | | | | | -5.3 | 1.3 |
| | Shola forest area | | | | | | | | | | | | | | | | | 0.3 | 0.2 |
| | Timber plantation area | | | | | | | | | | | | | | | | | -1.5 | 0.8 |
| | Proximity to stream (Topographical Wetness Index) | 0.2 | 0.0 | -0.1 | 0.0 | 0.0 | 0.1 | 0.0 | 0.0 | | | | | 0.3 | 0.0 | | | | |
