## Supplementary Table 2 for "Invasion blurs the biphasic mosaic, amplifies the ‘wicked problem’ through impacts on avian distributions in montane habitats"

Supplementary Table 2: Description of expected covariate effects on individual species' occupancy, abundance, and individual-level detectability. Abbreviations: −, negative effect; +, positive expected effect; NA, no expected effect, O, Occupancy; A, Abundance; D, Detectability.

| | | **Occupancy** | | | | | | | | | | | | | | | | | | | | | | | | | | |
|---|---|---|---|---|---|---|---|---|---|---|---|---|---|---|---|---|---|---|---|---|---|---|---|---|---|---|---|---|
| **Structural** | | **CUCE** | | | **EUAL** | | | **FINI** | | | **LATH** | | | **POHO** | | | **PYJO** | | | **SHKL** | | | **ZOPA** | | | **ANNI** | | |
| | **Variables** | O | A | D | O | A | D | O | A | D | O | A | D | O | A | D | O | A | D | O | A | D | O | A | D | O | A | D |
| | Canopy cover | + | + | - | + | + | - | | | | + | + | - | + | + | - | - | - | - | + | + | - | + | + | - | | | |
| | Ferns | + | + | + | | | | + | + | - | | | | | | | | | | | | | + | + | - | | | |
| | Leaf Litter | | | | | | | | | | + | + | - | | | | | | | | | | | | | | | |
| | Humidity (moss cover as a proxy) | + | + | - | + | + | - | + | + | - | + | + | - | + | + | - | | | | + | + | - | + | + | - | | | |
| | Total basal area | + | + | - | + | + | - | + | + | - | | | | + | + | - | - | - | - | | | | + | + | - | | | |
| | Foliage density b/w 0-2 m | + | + | - | + | + | - | + | + | - | | | | + | + | - | | | | | | | | | | | | |
| | Foliage density b/w 0-3 m | | | | | | | | | | | | | | | | | | | + | + | - | | | | | | |
| | Foliage density b/w 0-5 m | | | | | | | | | | + | + | - | + | + | - | + | + | - | | | | | | | | | |
| | Foliage density b/w 2-5 m | - | - | - | - | - | - | | | | | | | | | | | | | | | | + | + | - | | | |
| **Compositional** | Bamboo | | | | | | | + | + | - | | | | + | + | - | | | | | | | | | | | | |
| | Daphniphyllum | | | | + | + | + | | | | + | + | - | + | + | - | + | + | - | | | | | | | | | |
| | Lantana | | | | | | | | | | | | | | | | + | + | - | | | | | | | | | |
| | Rubus | | | | + | + | + | | | | + | + | - | + | + | - | + | + | - | | | | | | | | | |
| | Basal area of Shola trees | + | + | - | + | + | - | + | + | - | + | + | - | + | + | - | | | | | | | | | | | | |
| **Climatic** | Mean Temperature of Coldest Qtr | | | | - | - | - | - | - | - | | | | | | | | | | - | - | - | | | | - | - | - |

| | | | | | | | | | | | | | | | | | | | | | | | | | | | | |
|---|---|---|---|---|---|---|---|---|---|---|---|---|---|---|---|---|---|---|---|---|---|---|---|---|---|---|---|---|
| | Precipitation seasonality | - | - | - | | | | | | | - | - | - | | | | | | | - | - | - | | | | - | - | - |
| | Mean Precipitation of Warmest Qtr | | | | + | + | + | | | | | | | | | | | | | | | | | | | + | + | + |
| | Temp. Seasonality | - | - | - | | | | | | | - | - | - | - | - | - | + | + | + | | | | | | | - | - | - |
| **Landscape** | Elevation | + | + | + | + | + | + | + | + | + | + | + | + | + | + | + | | | | + | + | + | | | | | | |
| | Open human-modified area | | | | | | | | | | | | | | | | + | + | + | | | | | | | | | |
| | Productive area | + | + | + | + | + | + | | | | | | | | | | | | | | | | | | | + | + | + |
| | Distance from Shola edges (Natural wooded habitat) | - | - | - | - | - | - | - | - | - | - | - | - | - | - | - | + | + | + | - | - | - | - | - | - | | | |
| | Distance from Wooded habitat edges (Natural and/or transformed) | - | - | - | - | - | - | - | - | - | - | - | - | - | - | - | + | + | + | - | - | - | - | - | - | | | |
| | Distance from Grassland edges | | | | | | | | | | | | | | | | | | | | | | | | | - | - | - |
| | Distance from Open habitat edges (transformed) | | | | | | | | | | | | | | | | | | | | | | | | | - | - | - |
| | Shola forest area | | | | | | | | | | | | | | | | | | | | | | | | | + | + | + |
| | Timber plantation area | | | | | | | | | | | | | | | | | | | | | | | | | - | - | - |
| | Proximity to stream (Topographical Wetness Index) | + | + | + | + | + | + | | | | + | + | + | | | | | | | + | + | + | | | | | | |
| | Ruggedness | | | | | | | | | | | | | | | | | | | + | + | - | | | | | | |

| | | | | | | | | | | | | | | | | | | | | | | | | | | | | |
|---|---|---|---|---|---|---|---|---|---|---|---|---|---|---|---|---|---|---|---|---|---|---|---|---|---|---|---|---|
| **Visit-level variables** | Wind speed | NA | NA | - | NA | NA | - | NA | NA | - | NA | NA | - | NA | NA | - | NA | NA | - | NA | NA | - | NA | NA | - | NA | NA | NA |
| | Observer Experience | NA | NA | + | NA | NA | + | NA | NA | + | NA | NA | + | NA | NA | + | NA | NA | + | NA | NA | + | NA | NA | + | NA | NA | NA |
| **Ref.** | | 22,23 | | | 24,25 | | | 24,26 | | | 23,27,28 | | | 23,29 | | | 23,30 | | | 31–33 | | | 23,34 | | | | | |
