## Supplementary Table 3 for "Invasion blurs the biphasic mosaic, amplifies the ‘wicked problem’ through impacts on avian distributions in montane habitats"

Supplementary Table 3: Model selection for occupancy variation for the study birds. The table shows the four models with the lowest AIC values. The first tilde is followed by the covariate structure for detection, followed by the second tilde and the covariate structure for occurrence.

| Model_no | Model | AIC | Delta | AICwt |
|---|---|---|---|---|
| ANNI | | | | |
| ANNI.psi_occu70 | ~Observer + Day + Time + Distance from Edges of Open Areas+ Area of Shola forest within buffer around grid cell + Area of Timber plantation within buffer around grid cell + Temperature seasonality + Precipitation seasonality + Mean temperature of Coldest Quarter + Mean Precipitation of Warmest Quarter + offset(log(Area)) ~ Distance from Grassland Edges + Precipitation seasonality + Mean temperature of Coldest Quarter + Mean Precipitation of Warmest Quarter + Area of Shola forest within buffer around grid cell + Area of Timber plantation within buffer around grid cell | 832.96 | 0 | 0.17 |
| ANNI.psi_occu9 | ~Observer + Day + Time + Distance from Edges of Open Areas+ Area of Shola forest within buffer around grid cell + Area of Timber plantation within buffer around grid cell + Temperature seasonality + Precipitation seasonality + Mean temperature of Coldest Quarter + Mean Precipitation of Warmest Quarter + offset(log(Area)) ~ Distance from Grassland Edges + Temperature seasonality + Precipitation seasonality + Mean temperature of Coldest Quarter + Area of Shola forest within buffer around grid cell + Area of Timber plantation within buffer around grid cell | 833.03 | 0.08 | 0.17 |
| ANNI.psi_occuUniversal | ~Observer + Day + Time + Distance from Edges of Open Areas+ Area of Shola forest within buffer around grid cell + Area of Timber plantation within buffer around grid cell + Temperature seasonality + Precipitation seasonality + Mean temperature of Coldest Quarter + Mean Precipitation of Warmest Quarter + offset(log(Area)) ~ Distance from Grassland Edges + Temperature seasonality + Precipitation seasonality + Area of Shola forest within buffer around grid cell + Area of Timber plantation within buffer around grid cell + Mean temperature of Coldest Quarter + Mean Precipitation of Warmest Quarter | 833.40 | 0.45 | 0.14 |
| ANNI.psi_occu65 | ~Observer + Day + Time + Distance from Edges of Open Areas+ Area of Shola forest within buffer around grid cell + Area of Timber plantation within buffer around grid cell + Temperature seasonality + Precipitation seasonality + Mean temperature of Coldest Quarter + Mean Precipitation of Warmest Quarter + offset(log(Area)) ~ Distance from Grassland Edges + Temperature seasonality + Precipitation seasonality + Mean temperature of Coldest Quarter + Mean Precipitation of Warmest Quarter + Area of Shola forest within buffer around grid cell + Area of Timber plantation within buffer around grid cell | 833.40 | 0.45 | 0.14 |
| CUCE | | | | |
| CUCE.psi_woccuUniversal | ~Moss + Distance From Edges of Shola forest ~ Wetness + Elevation + Foliage density b/w 0-2 m + Foliage density b/w 2-5 m + Shola_ar + Moss + Ferns + Total.basal.area + Canopy + Distance From Edges of Wooded areas + Precipitation seasonality + Area of Productive Area Within buffer around Grid Cell | 3045.20 | 0.00 | 0.96 |
| CUCE.psi_woccu061 | ~Moss + Distance From Edges of Shola forest ~ Moss + Distance From Edges of Wooded areas | 3052.35 | 7.15 | 0.03 |
| CUCE.psi_occuUniversal | ~Moss + Distance From Edges of Shola forest ~ Wetness + Elevation + Foliage density b/w 0-2 m + Foliage density b/w 2-5 m | 3054.66 | 9.46 | 0.01 |

| | | | | |
|---|---|---|---|---|
| | + Shola_ar + Moss + Ferns + Total.basal.area + Canopy + Distance From Edges of Shola forest + Precipitation seasonality + Area of Productive Area Within buffer around Grid Cell | | | |
| CUCE.psi_occu095 | ~Moss + Distance From Edges of Shola forest ~ Ferns + Moss + Canopy | 3060.12 | 14.92 | 0.00 |
| EUAL | | | | |
| EUAL.psi_occu184 | ~Mean temperature of Coldest Quarter + Mean Precipitation of Warmest Quarter + Foliage density b/w 2-5 m + Canopy ~ Canopy + Distance From Edges of Shola forest + Rubus + Leaf_Litter | 2845.49 | 0.00 | 0.23 |
| EUAL.psi_occu262 | ~Mean temperature of Coldest Quarter + Mean Precipitation of Warmest Quarter + Foliage density b/w 2-5 m + Canopy ~ Canopy + Distance From Edges of Shola forest + Rubus + Leaf_Litter + Total.basal.area | 2847.16 | 1.67 | 0.10 |
| EUAL.psi_occu261 | ~Mean temperature of Coldest Quarter + Mean Precipitation of Warmest Quarter + Foliage density b/w 2-5 m + Canopy ~ Canopy + Distance From Edges of Shola forest + Rubus + Leaf_Litter + Shola_ar | 2847.28 | 1.79 | 0.09 |
| EUAL.psi_occu249 | ~Mean temperature of Coldest Quarter + Mean Precipitation of Warmest Quarter + Foliage density b/w 2-5 m + Canopy ~ Canopy + Distance From Edges of Shola forest + Moss + Rubus + Leaf_Litter | 2847.46 | 1.97 | 0.08 |
| FINI | | | | |
| FINI.psi_occuw205 | ~Wind + Observer + Foliage density b/w 0-2 m + Wetness + Total.basal.area + Distance From Edges of Wooded areas + Moss + Ferns + Shola_ar + Bamboo + Mean temperature of Coldest Quarter ~ Foliage density b/w 0-2 m + Distance From Edges of Wooded areas + Moss + Mean temperature of Coldest Quarter | 3026.54 | 0.00 | 0.57 |
| FINI.psi_occuUniversal | ~Wind + Observer + Foliage density b/w 0-2 m + Wetness + Total.basal.area + Distance From Edges of Wooded areas + Moss + Ferns + Shola_ar + Bamboo + Mean temperature of Coldest Quarter ~ Foliage density b/w 0-2 m + Wetness + Total.basal.area + Distance From Edges of Shola forest + Moss + Ferns + Shola_ar + Bamboo + Mean temperature of Coldest Quarter | 3029.60 | 3.05 | 0.12 |
| FINI.psi_occuw310 | ~Wind + Observer + Foliage density b/w 0-2 m + Wetness + Total.basal.area + Distance From Edges of Wooded areas + Moss + Ferns + Shola_ar + Bamboo + Mean temperature of Coldest Quarter ~ Foliage density b/w 0-2 m + Distance From Edges of Wooded areas + Moss + Ferns + Mean temperature of Coldest Quarter | 3030.63 | 4.09 | 0.07 |
| FINI.psi_occu310 | ~Wind + Observer + Foliage density b/w 0-2 m + Wetness + Total.basal.area + Distance From Edges of Wooded areas + Moss + Ferns + Shola_ar + Bamboo + Mean temperature of Coldest Quarter ~ Foliage density b/w 0-2 m + Distance From Edges of Shola forest + Moss + Ferns + Mean temperature of Coldest Quarter | 3030.69 | 4.15 | 0.07 |
| LATH | | | | |
| LATH.psi_occuUniversal | ~Wind + Observer + Foliage density b/w 0-5 m + Canopy + Wetness + Distance From Edges of Shola forest + Moss + Rubus + Daphniphyllum + Leaf_Litter + Shola_ar + Total.basal.area + Area of Productive Area Within buffer around Grid Cell + Precipitation seasonality + Temperature seasonality ~ Foliage density b/w 0-5 m + Canopy + Wetness + Distance From Edges of Shola forest + Moss + Rubus + Daphniphyllum + Leaf_Litter + Shola_ar + Total.basal.area + Area of Productive Area Within buffer around | 2832.52 | 0.00 | 0.64 |

| | | | | |
|---|---|---|---|---|
| | Grid Cell + Precipitation seasonality + Temperature seasonality | | | |
| LATH.psi_occuWUniversal | ~Wind + Observer + Foliage density b/w 0-5 m + Canopy + Wetness + Distance From Edges of Wooded areas + Moss + Rubus + Daphniphyllum + Leaf_Litter + Shola_ar + Total.basal.area + Area of Productive Area Within buffer around Grid Cell + Precipitation seasonality + Temperature seasonality ~ Foliage density b/w 0-5 m + Canopy + Wetness + Distance From Edges of Wooded areas + Moss + Rubus + Daphniphyllum + Leaf_Litter + Shola_ar + Total.basal.area + Area of Productive Area Within buffer around Grid Cell + Precipitation seasonality + Temperature seasonality | 2834.11 | 1.58 | 0.29 |
| LATH.psi_occu508 | ~Wind + Observer + Foliage density b/w 0-5 m + Canopy + Wetness + Distance From Edges of Shola forest + Moss + Rubus + Daphniphyllum + Leaf_Litter + Shola_ar + Total.basal.area + Area of Productive Area Within buffer around Grid Cell + Precipitation seasonality + Temperature seasonality ~ Precipitation seasonality + Temperature seasonality + Foliage density b/w 0-5 m + Canopy | 2838.27 | 5.75 | 0.04 |
| LATH.psi_occu488 | ~Wind + Observer + Foliage density b/w 0-5 m + Canopy + Wetness + Distance From Edges of Shola forest + Moss + Rubus + Daphniphyllum + Leaf_Litter + Shola_ar + Total.basal.area + Area of Productive Area Within buffer around Grid Cell + Precipitation seasonality + Temperature seasonality ~ Area of Productive Area Within buffer around Grid Cell + Precipitation seasonality + Temperature seasonality + Foliage density b/w 0-5 m | 2839.09 | 6.57 | 0.02 |
| POHO | | | | |
| POHO.psi_occuW073 | ~Foliage density b/w 0-5 m + Canopy + Total.basal.area + Temperature seasonality ~ Foliage density b/w 0-5 m + Moss + DistFrmWooded | 1236.46 | 0.00 | 0.02 |
| POHO.psi_occu401 | ~Foliage density b/w 0-5 m + Canopy + Total.basal.area + Temperature seasonality ~ Canopy + Moss + Temperature seasonality | 1236.60 | 0.15 | 0.02 |
| POHO.psi_occuW150 | ~Foliage density b/w 0-5 m + Canopy + Total.basal.area + Temperature seasonality ~ Foliage density b/w 0-5 m + Canopy + Moss + DistFrmWooded | 1236.86 | 0.40 | 0.02 |
| POHO.psi_occuW165 | ~Foliage density b/w 0-5 m + Canopy + Total.basal.area + Temperature seasonality ~ Foliage density b/w 0-5 m + Total.basal.area + Moss + DistFrmWooded | 1236.98 | 0.53 | 0.02 |
| PYJO | | | | |
| PYJO.psi_occu535 | ~Canopy + Rubus + Daphniphyllum + Lantana + Temperature seasonality ~ Canopy + Rubus + Daphniphyllum + Lantana + Temperature seasonality | 923.13 | 0.00 | 0.30 |
| PYJO.psi_occu296 | ~Canopy + Rubus + Daphniphyllum + Lantana + Temperature seasonality ~ Canopy + Rubus + Lantana + Temperature seasonality | 926.47 | 3.34 | 0.06 |
| PYJO.psi_occu542 | ~Canopy + Rubus + Daphniphyllum + Lantana + Temperature seasonality ~ Canopy + Rubus + Daphniphyllum + Temperature seasonality + Elevation | 926.88 | 3.76 | 0.05 |
| PYJO.psi_occu547 | ~Canopy + Rubus + Daphniphyllum + Lantana + Temperature seasonality ~ Canopy + Rubus + Lantana + Temperature seasonality + DistFrmShola | 926.90 | 3.77 | 0.05 |
| SHKL | | | | |

| | | | | |
|---|---|---|---|---|
| SHKL.psi_occuUniversal | ~Wetness + Distance From Edges of Shola forest + Precipitation seasonality + Mean temperature of Coldest Quarter ~ Foliage density b/w 0-3 m + Canopy + Wetness + Distance From Edges of Shola forest + Moss + Area of Productive Area Within buffer around Grid Cell + Precipitation seasonality + Mean temperature of Coldest Quarter | 2497.76 | 0.00 | 0.55 |
| SHKL.psi_woccuUniversal | ~Wetness + Distance From Edges of Shola forest + Precipitation seasonality + Mean temperature of Coldest Quarter ~ Foliage density b/w 0-3 m + Canopy + Wetness + Distance From Edges of Wooded areas + Moss + Area of Productive Area Within buffer around Grid Cell + Precipitation seasonality + Mean temperature of Coldest Quarter | 2499.38 | 1.61 | 0.25 |
| SHKL.psi_woccu180 | ~Wetness + Distance From Edges of Shola forest + Precipitation seasonality + Mean temperature of Coldest Quarter ~ Foliage density b/w 0-3 m + Moss + Distance From Edges of Wooded areas + Wetness + Area of Productive Area Within buffer around Grid Cell + Mean temperature of Coldest Quarter | 2502.44 | 4.68 | 0.05 |
| SHKL.psi_woccu168 | ~Wetness + Distance From Edges of Shola forest + Precipitation seasonality + Mean temperature of Coldest Quarter ~ Foliage density b/w 0-3 m + Canopy + Moss + Distance From Edges of Wooded areas + Area of Productive Area Within buffer around Grid Cell + Mean temperature of Coldest Quarter | 2503.25 | 5.49 | 0.04 |
| ZOPA | | | | |
| ZOPA.psi_occu017 | ~Foliage density b/w 2-5 m + Ferns + Total.basal.area + Precipitation seasonality ~ Canopy + Moss | 2841.54 | 0.00 | 0.05 |
| ZOPA.psi_occu002 | ~Foliage density b/w 2-5 m + Ferns + Total.basal.area + Precipitation seasonality ~ Canopy | 2842.06 | 0.51 | 0.04 |
| ZOPA.psi_occu045 | ~Foliage density b/w 2-5 m + Ferns + Total.basal.area + Precipitation seasonality ~ Canopy + Moss + Total.basal.area | 2842.13 | 0.58 | 0.04 |
| ZOPA.psi_occu018 | ~Foliage density b/w 2-5 m + Ferns + Total.basal.area + Precipitation seasonality ~ Canopy + Total.basal.area | 2842.96 | 1.41 | 0.03 |

Supplementary Table 4: Model selection for abundance variation for the study birds. The table shows the four models with the lowest AIC values. The first tilde is followed by the covariate structure for detection, followed by the second tilde and the covariate structure for abundance.

| Model_no | Model | AIC | delta | AICwt |
|---|---|---|---|---|
| CUCE | | | | |
| CUCE.psi_Universal | ~Canopy + Area of Productive Area Within buffer around Grid Cell ~ Wetness + Elevation + Foliage density b/w 0-2 m + Foliage density b/w 2-5 m + Shola_ar + Moss + Ferns + Total.basal.area + Canopy + Distance From Edges of Shola forest + Precipitation seasonality + Area of Productive Area Within buffer around Grid Cell + offset(log(Area)) | 5722.01 | 0.00 | 0.23 |
| CUCE.psi1_Universal | ~Wind + Area of Productive Area Within buffer around Grid Cell ~ Wetness + Elevation + Foliage density b/w 0-2 m + Foliage density b/w 2-5 m + Shola_ar + Moss + Ferns + Total.basal.area + Canopy + Distance From Edges of Shola forest + Precipitation seasonality + | 5722.29 | 0.28 | 0.20 |

| | | | | |
|---|---|---|---|---|
| | Area of Productive Area Within buffer around Grid Cell + offset(log(Area)) | | | |
| CUCE.psi_w Universal | ~Canopy + Area of Productive Area Within buffer around Grid Cell ~ Wetness + Elevation + Foliage density b/w 0-2 m + Foliage density b/w 2-5 m + Shola_ar + Moss + Ferns + Total.basal.area + Canopy + Distance From Edges of Wooded areas + Precipitation seasonality + Area of Productive Area Within buffer around Grid Cell + offset(log(Area)) | 5722.36 | 0.35 | 0.20 |
| CUCE.psi1_w Universal | ~Wind + Area of Productive Area Within buffer around Grid Cell ~ Wetness + Elevation + Foliage density b/w 0-2 m + Foliage density b/w 2-5 m + Shola_ar + Moss + Ferns + Total.basal.area + Canopy + Distance From Edges of Wooded areas + Precipitation seasonality + Area of Productive Area Within buffer around Grid Cell + offset(log(Area)) | 5722.72 | 0.71 | 0.16 |
| EUAL | | | | |
| EUAL.psi_Universal | ~Wetness + Mean Precipitation of Warmest Quarter + Foliage density b/w 0-2 m + Foliage density b/w 2-5 m ~ Foliage density b/w 0-2 m + Foliage density b/w 2-5 m + Canopy + Wetness + Distance From Edges of Shola forest + Moss + Rubus + Daphniphyllum + Leaf_Litter + Shola_ar + Total.basal.area + Area of Productive Area Within buffer around Grid Cell + Mean temperature of Coldest Quarter + Mean Precipitation of Warmest Quarter + offset(log(Area)) | 4198.67 | 0.00 | 0.91 |
| EUAL.psi_562 | ~Wetness + Mean Precipitation of Warmest Quarter + Foliage density b/w 0-2 m + Foliage density b/w 2-5 m ~ Distance From Edges of Shola forest + Mean temperature of Coldest Quarter + Foliage density b/w 0-2 m + Canopy + offset(log(Area)) | 4203.31 | 4.63 | 0.09 |
| EUAL.psi_463 | ~Wetness + Mean Precipitation of Warmest Quarter + Foliage density b/w 0-2 m + Foliage density b/w 2-5 m ~ Wetness + Distance From Edges of Shola forest + Mean temperature of Coldest Quarter + Canopy + offset(log(Area)) | 4210.52 | 11.85 | 0.00 |
| EUAL.psi_184 | ~Wetness + Mean Precipitation of Warmest Quarter + Foliage density b/w 0-2 m + Foliage density b/w 2-5 m ~ Canopy + Distance From Edges of Shola forest + Rubus + Leaf_Litter + offset(log(Area)) | 4212.88 | 14.21 | 0.00 |
| FINI | | | | |
| FINI.lambda_287 | ~Wind + Observer + Foliage density b/w 0-2 m + Wetness + Total.basal.area + Distance From Edges of Wooded areas + Moss + Ferns + Shola_ar + Bamboo + Mean temperature of Coldest Quarter ~ Foliage density b/w 0-2 m + Total.basal.area + Distance From Edges of Shola forest + Ferns + Mean temperature of Coldest Quarter + offset(log(Area)) | 4839.79 | 0.00 | 0.44 |
| FINI.lambda_Universal | ~Wind + Observer + Foliage density b/w 0-2 m + Wetness + Total.basal.area + Distance From Edges of Wooded areas + Moss + Ferns + Shola_ar + Bamboo + Mean temperature of Coldest Quarter ~ Foliage density b/w 0-2 m + Wetness + Total.basal.area + Distance From Edges of Shola forest + Moss + Ferns + Shola_ar + Bamboo + Mean temperature of Coldest Quarter + offset(log(Area)) | 4841.18 | 1.39 | 0.22 |
| FINI.lambda_wUniversal | ~Wind + Observer + Foliage density b/w 0-2 m + Wetness + Total.basal.area + Distance From Edges of Wooded areas + Moss + Ferns + Shola_ar + Bamboo + Mean temperature of Coldest Quarter ~ Foliage density b/w 0-2 m + Wetness + Total.basal.area + Distance From Edges of Wooded areas + Moss + Ferns + Shola_ar + Bamboo + Mean temperature of Coldest Quarter + offset(log(Area)) | 4842.84 | 3.05 | 0.10 |

| | | | | |
|---|---|---|---|---|
| FINI.lambda_285 | ~Wind + Observer + Foliage density b/w 0-2 m + Wetness + Total.basal.area + Distance From Edges of Wooded areas + Moss + Ferns + Shola_ar + Bamboo + Mean temperature of Coldest Quarter ~ Foliage density b/w 0-2 m + Total.basal.area + Distance From Edges of Shola forest + Moss + Mean temperature of Coldest Quarter + offset(log(Area)) | 4843.63 | 3.84 | 0.06 |
| | | LATH | | |
| LATH.psi_wUniversal | ~Wind + Observer + Foliage density b/w 0-5 m + Canopy + Wetness + Distance From Edges of Wooded areas + Moss + Rubus + Daphniphyllum + Leaf_Litter + Shola_ar + Total.basal.area + Area of Productive Area Within buffer around Grid Cell + Precipitation seasonality + Temperature seasonality ~ Foliage density b/w 0-5 m + Canopy + Wetness + Distance From Edges of Wooded areas + Moss + Rubus + Daphniphyllum + Leaf_Litter + Shola_ar + Total.basal.area + Area of Productive Area Within buffer around Grid Cell + Precipitation seasonality + Temperature seasonality + offset(log(Area)) | 5744.76 | 0.00 | 0.89 |
| LATH.psi_w478 | ~Wind + Observer + Foliage density b/w 0-5 m + Canopy + Wetness + Distance From Edges of Wooded areas + Moss + Rubus + Daphniphyllum + Leaf_Litter + Shola_ar + Total.basal.area + Area of Productive Area Within buffer around Grid Cell + Precipitation seasonality + Temperature seasonality ~ Distance From Edges of Wooded areas + Temperature seasonality + Foliage density b/w 0-5 m + Canopy + offset(log(Area)) | 5748.91 | 4.15 | 0.11 |
| LATH.psi_Universal | ~Wind + Observer + Foliage density b/w 0-5 m + Canopy + Wetness + Distance From Edges of Wooded areas + Moss + Rubus + Daphniphyllum + Leaf_Litter + Shola_ar + Total.basal.area + Area of Productive Area Within buffer around Grid Cell + Precipitation seasonality + Temperature seasonality ~ Foliage density b/w 0-5 m + Canopy + Wetness + Distance From Edges of Shola forest + Moss + Rubus + Daphniphyllum + Leaf_Litter + Shola_ar + Total.basal.area + Area of Productive Area Within buffer around Grid Cell + Precipitation seasonality + Temperature seasonality + offset(log(Area)) | 5757.51 | 12.75 | 0.00 |
| LATH.psi_478 | ~Wind + Observer + Foliage density b/w 0-5 m + Canopy + Wetness + Distance From Edges of Wooded areas + Moss + Rubus + Daphniphyllum + Leaf_Litter + Shola_ar + Total.basal.area + Area of Productive Area Within buffer around Grid Cell + Precipitation seasonality + Temperature seasonality ~ Distance From Edges of Shola forest + Temperature seasonality + Foliage density b/w 0-5 m + Canopy + offset(log(Area)) | 5764.25 | 19.48 | 0.00 |
| | | POHO | | |
| POHO.psi_w406 | ~Foliage density b/w 0-5 m + Daphniphyllum + DistFrmWooded + Temperature seasonality ~ Canopy + DistFrmWooded + Temperature seasonality + offset(log(Area)) | 2295.28 | 0.00 | 0.10 |
| POHO.psi_404 | ~Foliage density b/w 0-5 m + Daphniphyllum + DistFrmWooded + Temperature seasonality ~ Canopy + Daphniphyllum + Temperature seasonality + offset(log(Area)) | 2296.30 | 1.02 | 0.06 |
| POHO.psi1_404 | ~Foliage density b/w 0-5 m + Total.basal.area + Daphniphyllum + Temperature seasonality ~ Canopy + Daphniphyllum + Temperature seasonality + offset(log(Area)) | 2296.39 | 1.11 | 0.06 |
| POHO.psi1_406 | ~Foliage density b/w 0-5 m + Total.basal.area + Daphniphyllum + Temperature seasonality ~ Canopy + DistFrmShola + Temperature seasonality + offset(log(Area)) | 2296.70 | 1.42 | 0.05 |
| | | PYJO | | |

| | | | | |
|---|---|---|---|---|
| PYJO.lambda_w_Universal | ~Canopy + Total.basal.area + Daphniphyllum + Temperature seasonality + DistFrmWooded ~ Foliage density b/w 0-5 m + Canopy + Total.basal.area + Rubus + Daphniphyllum + Lantana + DistFrmWooded + Area of Open Area Within buffer around Grid Cell + Temperature seasonality + Elevation + offset(log(Area)) | 4870.04 | 0.00 | 0.46 |
| PYJO.lambda_w_iUniversal | ~Foliage density b/w 0-5 m + Canopy + Total.basal.area + Daphniphyllum + DistFrmWooded ~ Foliage density b/w 0-5 m + Canopy + Total.basal.area + Rubus + Daphniphyllum + Lantana + DistFrmWooded + Area of Open Area Within buffer around Grid Cell + Temperature seasonality + Elevation + offset(log(Area)) | 4870.59 | 0.55 | 0.35 |
| PYJO.lambda_Universal | ~Canopy + Total.basal.area + Daphniphyllum + Temperature seasonality + DistFrmWooded ~ Foliage density b/w 0-5 m + Canopy + Total.basal.area + Rubus + Daphniphyllum + Lantana + DistFrmShola + Area of Open Area Within buffer around Grid Cell + Temperature seasonality + Elevation + offset(log(Area)) | 4872.82 | 2.79 | 0.11 |
| PYJO.lambda_iUniversal | ~Foliage density b/w 0-5 m + Canopy + Total.basal.area + Daphniphyllum + DistFrmWooded ~ Foliage density b/w 0-5 m + Canopy + Total.basal.area + Rubus + Daphniphyllum + Lantana + DistFrmShola + Area of Open Area Within buffer around Grid Cell + Temperature seasonality + Elevation + offset(log(Area)) | 4874.52 | 4.48 | 0.05 |
| SHKL | | | | |
| SHKL.lambda_Universal | ~Canopy + Wetness + Area of Productive Area Within buffer around Grid Cell + Mean temperature of Coldest Quarter ~ Foliage density b/w 0-3 m + Canopy + Wetness + Distance From Edges of Shola forest + Moss + Area of Productive Area Within buffer around Grid Cell + Precipitation seasonality + Mean temperature of Coldest Quarter + offset(log(Area)) | 3466.44 | 0.00 | 0.82 |
| SHKL.lambda_188 | ~Canopy + Wetness + Area of Productive Area Within buffer around Grid Cell + Mean temperature of Coldest Quarter ~ Canopy + Moss + Distance From Edges of Shola forest + Area of Productive Area Within buffer around Grid Cell + Precipitation seasonality + Mean temperature of Coldest Quarter + offset(log(Area)) | 3469.76 | 3.32 | 0.16 |
| SHKL.lambda_168 | ~Canopy + Wetness + Area of Productive Area Within buffer around Grid Cell + Mean temperature of Coldest Quarter ~ Foliage density b/w 0-3 m + Canopy + Moss + Distance From Edges of Shola forest + Area of Productive Area Within buffer around Grid Cell + Mean temperature of Coldest Quarter + offset(log(Area)) | 3475.15 | 8.71 | 0.01 |
| SHKL.lambda_177 | ~Canopy + Wetness + Area of Productive Area Within buffer around Grid Cell + Mean temperature of Coldest Quarter ~ Foliage density b/w 0-3 m + Canopy + Distance From Edges of Shola forest + Area of Productive Area Within buffer around Grid Cell + Precipitation seasonality + Mean temperature of Coldest Quarter + offset(log(Area)) | 3476.32 | 9.88 | 0.01 |
| ZOPA | | | | |
| ZOPA.psi_132 | ~Wind + Observer + Foliage density b/w 2-5 m + Canopy + Ferns + Moss + Total.basal.area + Area of Productive Area Within buffer around Grid Cell + Precipitation seasonality + Temperature seasonality + Distance From Edges of Wooded areas ~ Canopy + Ferns + Moss + Temperature seasonality + offset(log(Area)) | 11055.47 | 0.00 | 0.19 |
| ZOPA.psi_131 | ~Wind + Observer + Foliage density b/w 2-5 m + Canopy + Ferns + Moss + Total.basal.area + Area of Productive Area Within buffer around Grid Cell + Precipitation seasonality + Temperature seasonality + Distance From Edges of Wooded areas ~ Canopy + Ferns + Moss + Precipitation seasonality + offset(log(Area)) | 11055.52 | 0.05 | 0.18 |

| | | | | |
|---|---|---|---|---|
| ZOPA.psi_Universal | ~Wind + Observer + Foliage density b/w 2-5 m + Canopy + Ferns + Moss + Total.basal.area + Area of Productive Area Within buffer around Grid Cell + Precipitation seasonality + Temperature seasonality + Distance From Edges of Wooded areas ~ Foliage density b/w 2-5 m + Canopy + Ferns + Moss + Total.basal.area + Area of Productive Area Within buffer around Grid Cell + Precipitation seasonality + Temperature seasonality + Distance From Edges of Shola forest + offset(log(Area)) | 11055.65 | 0.18 | 0.17 |
| ZOPA.psi_wUniversal | ~Wind + Observer + Foliage density b/w 2-5 m + Canopy + Ferns + Moss + Total.basal.area + Area of Productive Area Within buffer around Grid Cell + Precipitation seasonality + Temperature seasonality + Distance From Edges of Wooded areas ~ Foliage density b/w 2-5 m + Canopy + Ferns + Moss + Total.basal.area + Area of Productive Area Within buffer around Grid Cell + Precipitation seasonality + Temperature seasonality + Distance From Edges of Wooded areas + offset(log(Area)) | 11056.41 | 0.94 | 0.12 |
