## Supplementary Fig 1 for "Invasion blurs the biphasic mosaic, amplifies the ‘wicked problem’ through impacts on avian distributions in montane habitats"

SUPPLEMENTARY FIGURES:

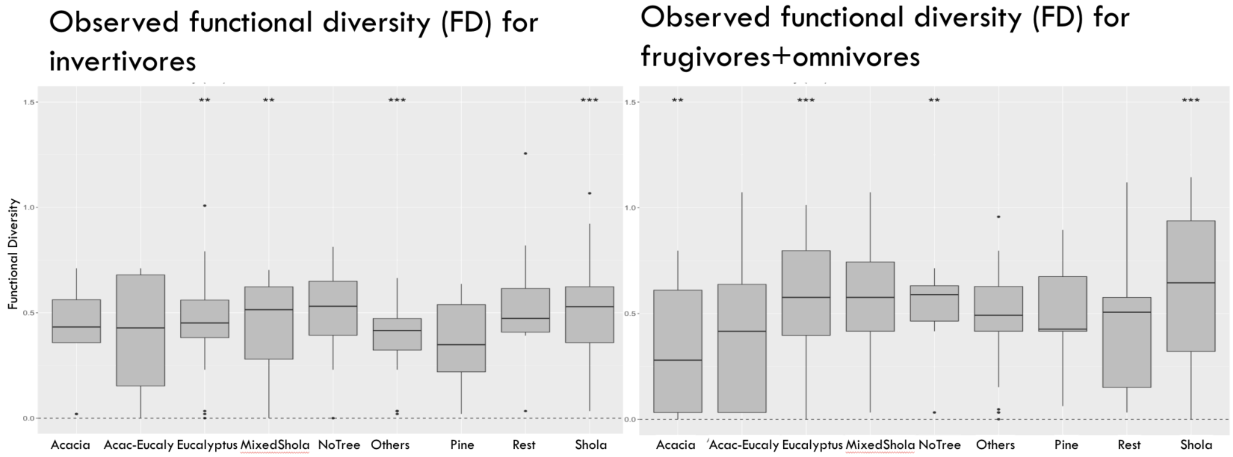


Supplementary Fig 1: (A): 169 avian communities across nine habitats. Asterisks indicate that the observed FD was significantly different from the null expectation (*<0.05, **<0.01, ***<0.001). Wooded habitats: Acacia (stands of *Acacia mearnsii*), Acacia-Eucalyptus (stands of *Acacia mearnsii* mixed with *Eucalyptus spp.*), Eucalyptus (stands of *Eucalyptus spp.*), Mixed Shola (stands with shola trees mixed with other non-native trees), Pine (stands of *Pinus spp.*), Shola (stands with only Shola trees), Rest (Mix of orchard trees). Open habitats: No trees (Sites with no trees). All statistical results are from two-tailed Wilcoxon signed-ranks tests.
